# Collagen of ancient bones gives an indication of endogenous DNA preservation based on the next generation sequencing technology

**DOI:** 10.1101/2023.05.02.538509

**Authors:** Yuka Nakamura, Daisuke Waku, Yoshiki Wakiyama, Yusuke Watanabe, Kae Koganebuchi, Tomohito Nagaoka, Kazuaki Hirata, Jun Ohashi, Ryuzaburo Takahashi, Minoru Yoneda, Hiroki Oota

## Abstract

Ancient genome analysis has become an indispensable tool in studies of human population history and evolution after the breakthrough of whole genome sequencing technology. Meanwhile, the problem has been not resolved that ancient genome cannot be analyzed without crushing non-small pieces of precious specimens; in spite of that, in many cases, there is no DNA remaining sufficiently in the piece of samples for obtaining the whole genome sequences. In previous studies, therefore, a couple of indicators (e.g., racemization ratios) have been proposed to estimate the endogenous DNA in ancient samples. However, these studies have used polymerase chain reaction (PCR) to test whether endogenous DNA is remaining, which has been inadequate because the success or failure of PCR amplification does not necessarily reflect DNA remaining. To assess the amount of endogenous DNA, we use the ratios of reads generated by the next-generation sequencer (NGS) mapped to the human reference genome sequence. We investigate 40 Jomon remains and find a significant association between the collagen residual ratios (CRRs) in rib bones and the mapping ratios (MRs). Because the weight of bone required to measure collagen residual is much less than that required to obtain DNA needed for NGS analysis, which is always necessary for dating, the collagen in ribs can be a good indicator for successful ancient genome analyses.

## Introduction

DNA extracted from ancient biological remains provides important information on evolution and diversity of organisms in the past. In the early days, ancient DNA sequence analyses relied on the dideoxy (Sanger) method through PCR amplification (Pääbo et al., 1988; Hagelberg et al., 1989; Pääbo et al., 1989; Lawlor et al., 1991; Kurosaki et al., 1993; Oota et al., 1995; Shinoda and Kanai, 1999). The PCR-based Sanger sequencing, however, gave us only a limited piece of genome information. The whole genome sequencing became more quickly and cheaper with the advent of NGS in the 21st century, and the technology was applied to ancient genome analyses for archaic hominins (e.g. Green et al., 2006, 2010; Krause et al. 2010; Reich et al. 2010; Meyer et al., 2012; Prufer et al. 2014) and modern humans (e.g. Fu et al., 2014; Rasmussen et al., 2010; Kanzawa-Kiriyama et al., 2017, 2019; McColl et al., 2018; Gakuhari et al., 2020). These research has provided us with enormous information about relationships with archaic hominins and peopling history of modern humans spread from Africa to other continents.

Despite the technological advance in DNA sequencing, genome analyses of paleo-biological subjects are fraught with difficulties: ancient DNA undergoes chemical modifications in the postmortem of the organisms, fragmenting and reducing the number of molecules. Soft and hard tissues of organisms are degraded by bacteria and fungi that enter through the surrounding soil in which the remains are buried, and some environmental factors, such as pH, temperature, exposure to water (Lindahl, 1993; Hedges and Millard, 1995), and humidity (Pinhasi et al., 2015) affect the rate of DNA molecule degradation. Because the DNA strand is fragmented and the number of DNA molecules is reduced dramatically, through the unavoidable process mentioned above, the ancient samples to be analyzed do not always contain sufficient amounts of DNA for genome sequencing. This difficulty concerns the other difficulty in selecting samples for ancient genome analysis. Even if valuable ancient samples are crushed for DNA extraction, they could be wasted if DNAs are not available.

For these reasons, a non-crushing or a nearly non-crushing indicator for estimating DNA preservation has been sought in previous studies for minimizing the damage of valuable biological remains to be analyzed. Initially, the rate of racemization of amino acids was used to assess a state of DNA preservation in ancient biological remains (Poinar et al., 1996). Nextly, the collagen residual ratios and crystallinity of hydroxyapatite in the specimens were reported as such indicators (Götherström et al., 2002). Both the racemization rate and the collagen residual rate gave certain criteria, but whether DNA remained or not was determined on the basis of whether PCR amplification could or could not be achieved successfully. It is not ruled out the possibilities of unsuccessful amplification due to the principle of the PCR method, and of false-positive amplification with contamination. Hence, a non-PCR based method for assessing a state of DNA preservation in ancient specimens has been required to develop.

Here we propose an NGS-based method to obtain a more quantitative indicator for DNA preservation exclusively in human skeletal remains. We consider the rate at which reads from shotgun sequencing map to the human reference sequence as the percentage of DNA remaining in the ancient sample. We first compared whether environmental factors (in- /out-shell-stratum and calibrated radiocarbon dates or the molecular factor (collagen residual ratio: CRR) correlated with the mapping ratio (MR). The results showed that collagen residual ratios, as expected from previous studies, correlated best with mapping ratios, though no association was shown in both environmental factors. The finding was that the association was statistically significant, especially in those of rib bones. This could be useful to assess the amount of DNA remaining and give a hint to develop a method avoiding unnecessary crushing of precious ancient human bones.

## Materials & Methods

### Archaeological samples

We examined 40 individuals excavated from three shell-mound sites (15 from Gionbara [GB], 7 Saihiro [SH], and 18 from Kikumatenaga [KT]) where locate in Ichihara City, Chiba Prefecture, and close to each other within 10 km (Figure 1). Based on the chronological age of earthenware, the Gionbara and Saihiro sites were assigned to the late to last Jomon period (4,400 – 2,400 years ago), whereas the Kikumatenaga site was assigned to the late Jomon period (3,200 – 2,400 years ago).

**Figure 1.**
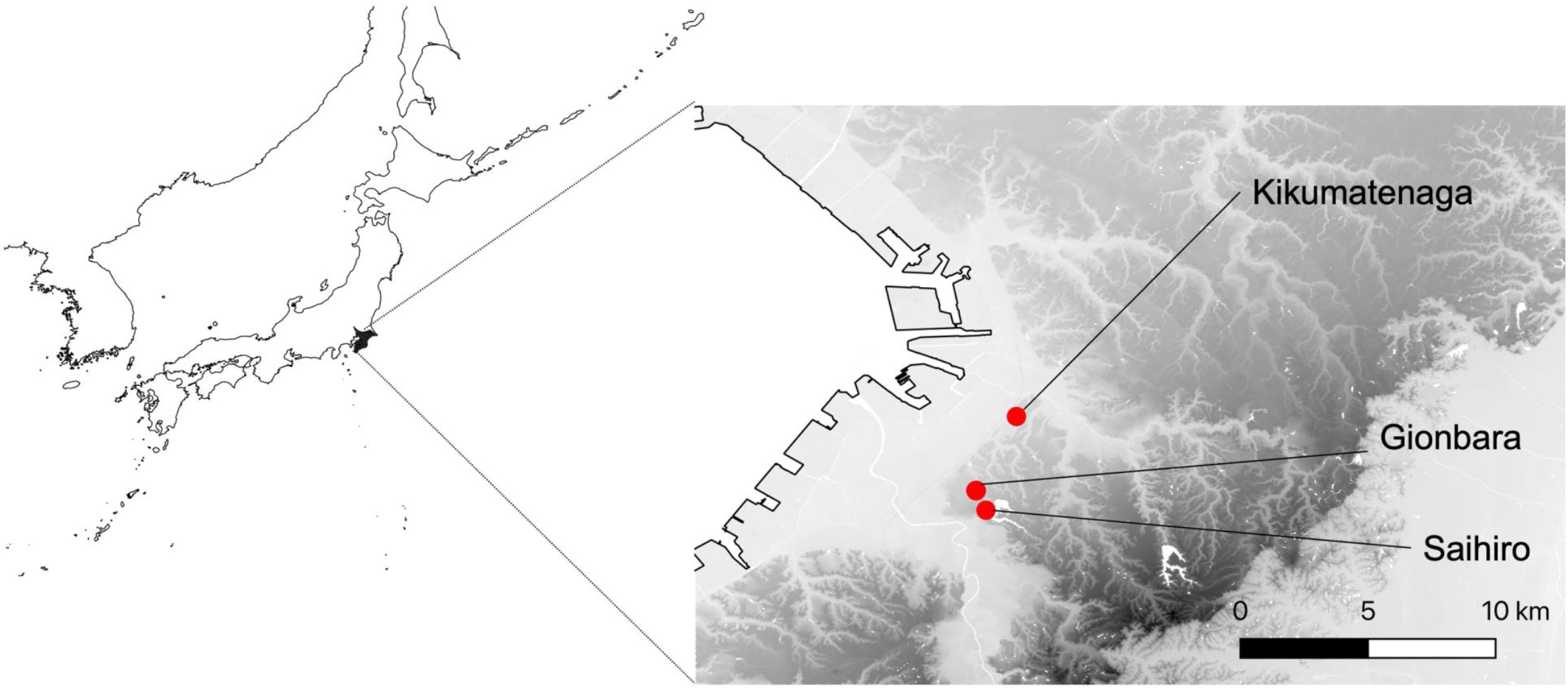
The geographical location of three archaeological sites (Gionbara, Kikumatenaga, and Saihiro shell mounds)

Regarding the SH and KT sites, all human skeletal remains were excavated from the shell stratum. Meanwhile, regarding the GB site, four individuals were excavated from the shell stratum; ten were excavated from the non-shell layer; one was unknown. (Oshizawa, 1999; Kondo, 1987; Sakurai, 2005). Whether a shell or non-shell stratum affects the pH of the surrounding sediments.

### Gelatin extraction and C14 dating

Each sample was subjected to collagen extraction from one different part (22 ribs, 11 skulls; 5 limbs; 1 ilium). Gelatin extraction was based on the protocols in previous studies (Longin, 1971; Yoneda et al., 2002). Bone samples were cleaned by sandblasting and ultrasonic cleaning for 10 min. They were demineralized for 40 h with 0.4 M HCl and neutralized for 23 h 50 min with pure water. After first neutralization, samples were treated with 0.1 M NaOH for 1 h and neutralized for 3 h 30 min. After the second neutralization, samples were gelatinized for 41 h at 90 °C with 0.0001 M HCl (pH 4.0). Suction filtration was done by Whatman GF/F. Filtrated samples were freeze-dried, and the amount of gelatin was weighed. Henceforth, this weight was considered as collagen content. Radiocarbon dating was measured by Accelerator Mass Spectrometry (AMS) at The University Museum at The University of Tokyo. The dates were referenced to IntCal20 and Marine 20 (Reimer et al. 2020, Heaton et al. 2020) calibrated ages were estimated using OxCal 4.4 software (Bronk Ramsey, 2009). The marine reservoir effect, in which the apparent radiocarbon dates become older due to sea food exploitation, was corrected by evaluating the effect of marine carbon based on δ^13^C. Using the mean values (−22.6‰ and -10.9‰) of land mammal and marine fish bones excavated from shell middens in Chiba Prefecture as endmembers, we estimated the marine contribution and mixed IntCal20 and Marine20, assuming an error of 5%. A regional correction value (ΔR) for Tokyo Bay was adapted to Marine20 (−98 ± 37 years) based on a shell collected in 1882 AD (534 ± 36 BP; Yoshida et al. 2010). The median of the probability distribution of dating was used as the estimated age.

### DNA extraction

Petrous bones from 40 individuals were used in this study. DNA was extracted following the protocols shown in previous studies (Gakuhari et al., 2020; Gamba et al., 2014). The petrous bone samples were cut by sterile disc cutters and drills to about 100 mg. The bone pieces in 5 ml tube of DNA Lobind were treated pre-digestion for 15 minutes at 900 rpm in a Thermomixer (Eppendorf, Hamburg, Germany) with the lysis buffer (2 ml) containing 20 mM Tris HCl (pH 7.4), 0.7% N-Lauroylsarcosine sodium salt solution, 0.5 M EDTA (pH 8.0), 1.2 U/ml recombinant Proteinase K. After the supernatant was transferred, the lysis buffer was added to the sample. It was placed into Thermomixer at 900 rpm and 60 °C for overnight (>16 h). After digestion, the tube was centrifuged for 10 min at 3500 rpm. The supernatant diluted with 10 mM TE buffer was added to the Filter Unit (Amicon Ultra 4, 10 kDa), and centrifuged for 30 min at 3500 rpm, or until a final volume of 100 μl was reached. The final volume was added to a Qiagen Mini elute column with a 2 ml tube and purified according to the instruction. The change of the instruction was adding Tween 20 to EB buffer pre-heated 60 °C.

All experiments were performed in the clean room exclusively for ancient genome experiments in the Department of Biological Sciences, Graduate School of Science, University of Tokyo. Before and after use, the room was illuminated with UV light. The clean room in the working state was kept at a positive pressure and air without modern DNA was supplied through a HEPA filter. The researchers always wore a body suit, face mask, and gloves. The tools and bone surfaces were exposed to UV.

### Library construction

The 40 libraries were constructed with NEBNext® Ultra II DNA Library Prep Kit for Illumina® (New England Biolabs, Ipswich, MA) according to the instruction. The change in the instruction was adaptor dilution and the amount of beads volume. The adaptor dilution was changed from 25-Fold to 10-Fold. The amount of 1st beads addition was changed from 40 μl to 90 μl. Similarly, the amount of 2nd beads addition was changed from 20 μl to 90 μl. The concentration of DNA extracts and libraries was measured by Qubit (Thermofisher, Waltham, MA). The fragment length of DNA extracts and libraries was checked by Bioanalyzer (Agilent Technologies, Santa Clara, CA). The sequencing was carried out in MiSeq (Illumina, San Diego, CA). The libraries were sequenced on a flow cell using the MiSeq Reagent Kit v3 150 cycles in paired-end.

### Raw sequencing data processing

The fastq files generated by MiSeq were processed by the following pipeline. AdaptorRemoval v2.2.2 (Schubert et al., 2016) was used to trim ambiguous bases at termini (-trimns), low-quality bases at termini (-trimqualities), short leads (-minlength 35) and to combine paired-end reads into consensus sequence (-collapse). BWA v0.7.16 (Li and Durbin, 2009) was used to map combined reads to human genome reference hg19. CleanSam in Picard Tools v2.21.8 was used to soft-clip beyond-end-of reference alignments and to set mapping quality to 0 for unmapped reads. Duplicate reads were removed by MarkDuplicates in Picard Tools (https://broadinstitute.github.io/picard/). The mapping ratio (the ratio of mapped reads divided by the number of reads) was calculated by flagstat in SAMtools v1.10 (Danecek et al., 2021). The substitution pattern of DNA molecules in libraries was checked by mapDamage2 v2.2.0 (Jónsson et al., 2013).

### Statistical analysis

The Mann-Whitney U test was used to compare mapping ratios between bones in the shell layers and those not in the shell layers. Linear regression analysis was conducted to examine the association of carbon 14 dates with mapping ratios, and the association of collagen residual ratios with mapping ratios. Furthermore, the association between collagen residual ratios and mapping ratios was examined separately for the skulls and the ribs. The collagen residual ratio was defined as gelatin weight divided by bone weight. The association of collagen residual ratios with mapping ratios was also assessed by a linear model with two dummy variables representing three remains (SH, KT, or GB). In the analysis, GB was used as the reference. The R package “coin” was used to carry out the Mann-Whitney U test. The R package “tidyverse” and “forcats” were used to carry out the regression analysis.

## Result

We summarize the mapping ratios (MRs), collagen residual ratios (CRRs), and carbon-to-nitrogen (CN) ratios of all the samples (Table 1). The MR represents a proxy of the rate of endogenous DNA. For instance, we obtained 2.55 ng/ul of DNA and 0.01% of mapping ratio from GB2-63, which represents almost all DNA extracted from this sample was from non-human organisms. Among 40 samples, the best MR was obtained at 62.97% in KT69. As observed in previous studies on ancient DNA, the MRs varied from each sample; those ranges were 0.01 – 44.52%, 9.03 – 48.64%, and 0.01 – 62.97% in GB, SH, KT, respectively, showing typical patterns of deamination (Supplementary Fig. 1). Radiocarbon dating was measured for 39 samples (one out of 40 was failed to be measured). The CRR was 0.5 - 9.6%, and the CN ratio was 2.9 to 3.6, the range of which indicates the collagen was not denatured (DeNiro, 1985), except for three out of 39 samples. Because there was no relationship between the MR and the CN ratio, we used all the CRRs (including the three samples) in the subsequent analyses. We successfully measured radiocarbon dating for 35 samples but failed for four out of 39 samples because their gelatins were not extracted. The estimated dating was 5076 – 3,424 calBP. The numbers of samples that were used for each testing an association between environmental and/or molecular factors and mapping ratios were summarized in Table 2.

**Table 1.**
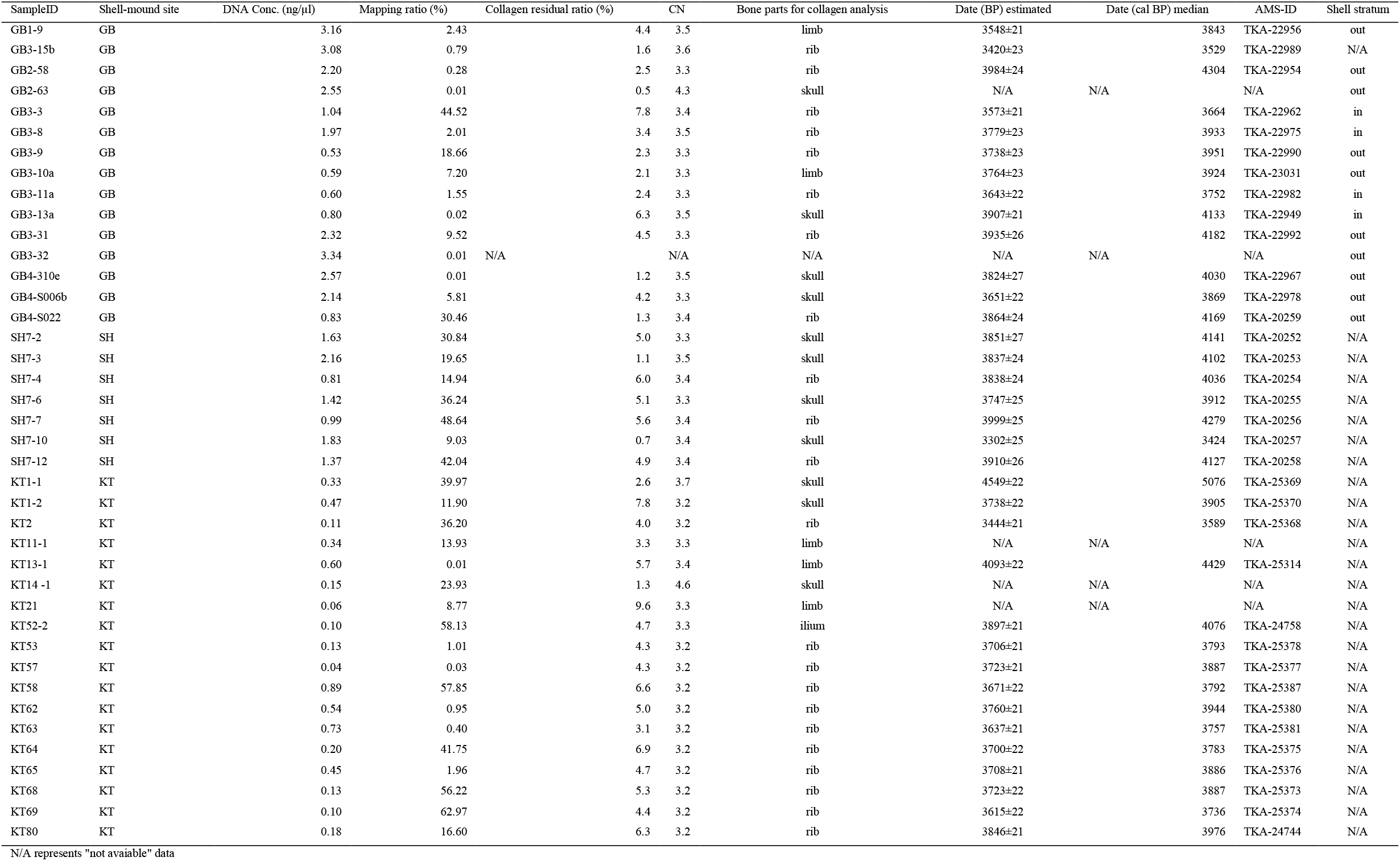
Summary of the measurments in DNA and collagen

**Table 2.** The numbers of the samples for testing correlation with the factors

|  | Shell stratum | Dating | Collagen | Total |
| --- | --- | --- | --- | --- |
| Gionbara [GB] | 14 | 13 | 14 | 15 |
| Saihiro [SH] | N/A | 7 | 7 | 7 |
| Kikumatenaga [KT] | N/A | 15 | 18 | 18 |
| Total | 14 | 35 | 39 | 40 |

First, we examined an association between the environmental factors and the MRs. The previous studies reported that the bones were in relatively good condition when the remains were buried in a stratum that was slightly alkaline (Gordon and Buikstra, 1981; Stephan, 1997). In general, the sedimental pH of in-shell-stratum is higher than that of out-shell-stratum. We expected, therefore, DNA extracted from the bones in the shell-stratum could be in a better condition than that out the shell-stratum. Among the three sites, only the GB site was available for the comparative test, because all the skeletal remains in the SH and KT sites were excavated from the in-shell-stratum (Table 2). Four out of 14 samples were excavated from the in-shell-stratum and the others were from the out-shell-stratum in the GB site. We found, however, no significant difference between the in- and out-shell-stratum in MRs (Mann-Whitney U-test, *p* = 0.887) (Figure 2). Similarly, no significant association was found between the dating and the MRs (*p* = 0.656) (Figure 3).

**Figure 2.**
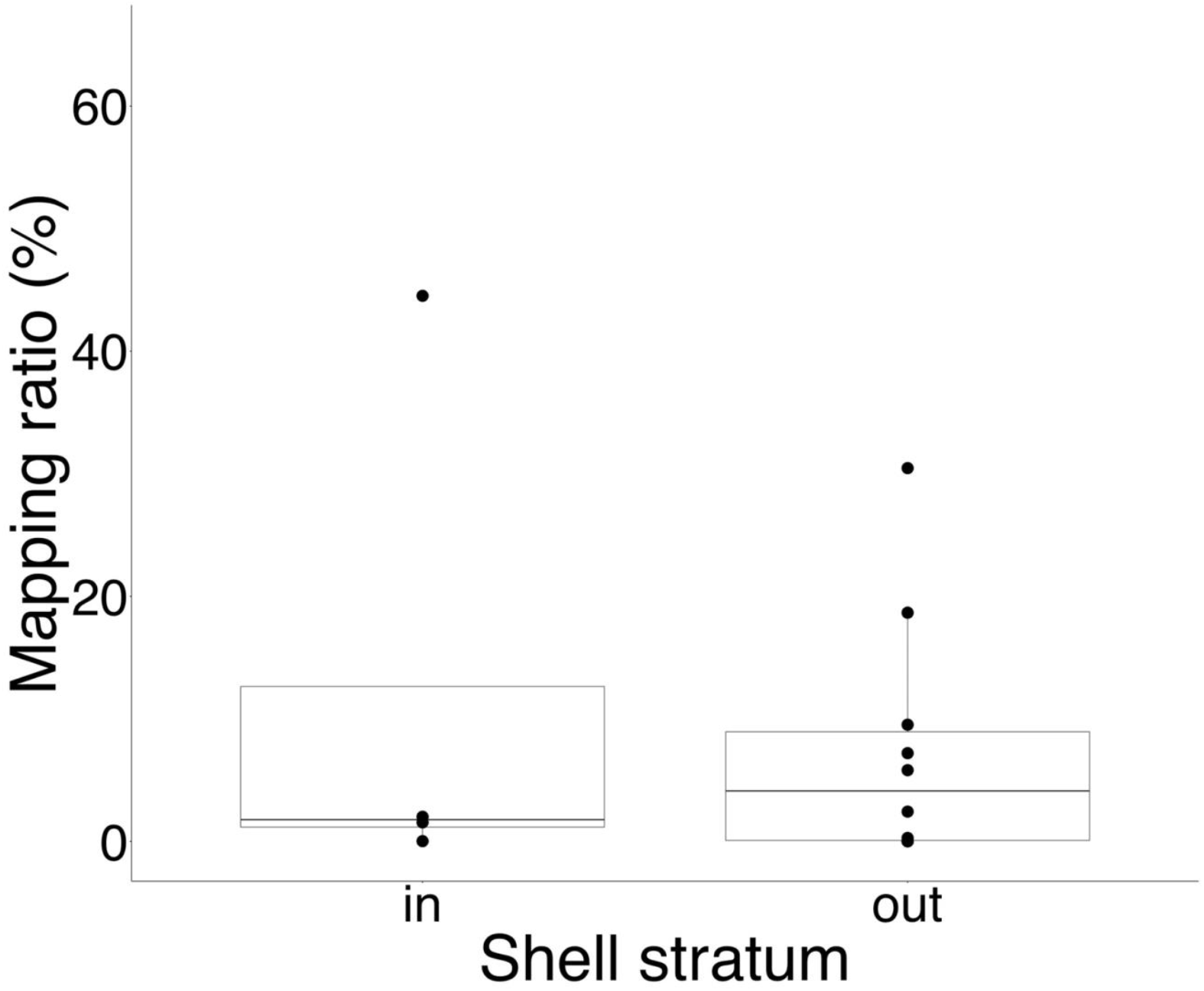
The association between the environmental factors (in- /out-shell-stratum) and the MRs. Four individuals were excavated from the shell layer; ten were excavated from the non-shell layer. There was no significant difference between them (*p* = 0.887).

**Figure 3.**
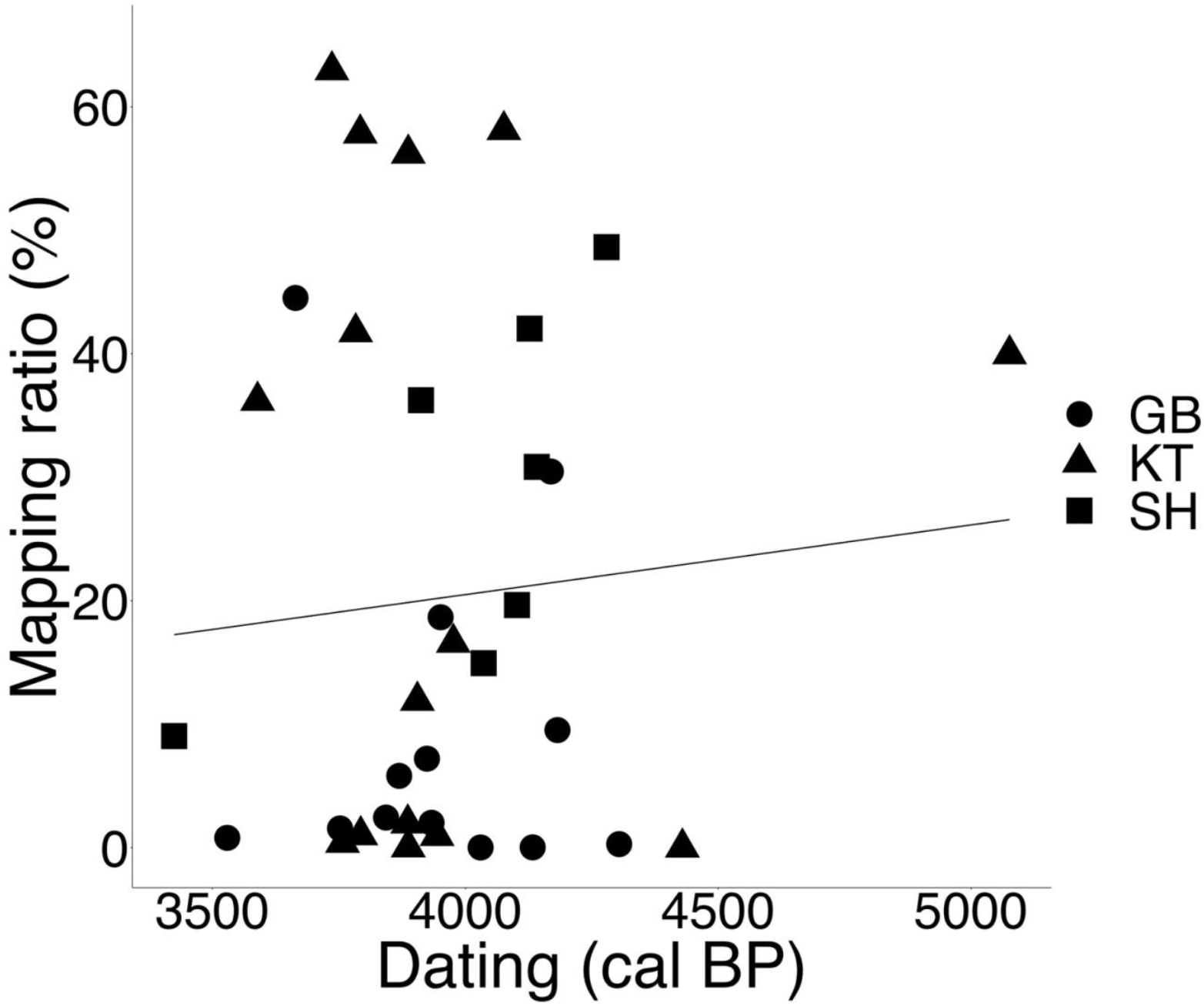
The association between the environmental factors (dating) and the MRs in the three Jomon sites. There was no association between them (*p* = 0.651). Circles, triangles, and squares represent the Gionbara shell-mound site (n = 13), the Kikumatenaga shell-mound site (n = 15), the Saihiro shell-mound site (n = 7), respectively.

Second, we tested an association between a molecular factor, CRR, and MR. The gelatins were extracted from the 39 bone samples and the CRRs were calculated as a gelatin weight divided by a bone weight. Overall, there was no significant association between the 39 CRRs from all archaeological sites and their MRs (*p* = 0.091) (Figure 4a). When the association was examined for each site, no significant association was found between them (*p* = 0.160 for GB; *p* = 0.146 for SH; *p* = 0.939 for KT). Regarding the gelatin extraction for estimating the CRRs, we used different parts of bones for each sample (see Table 1). The relationship between the CRRs and the MRs of different bone parts (ilium, limb, rib, and skull) was shown in Figure 4a. Among the four parts of bones, the statistical tests were performed only for skulls (n = 11) and for ribs (n = 22), since the numbers of samples were small for limb (n = 5) and ilium (n = 1). No significant association was found between the CRRs and the MRs for the skulls (*p* = 0.745) (Figure 4b), while a significant association was detected between CRRs and MRs for ribs (*p* = 0.020) (Figure 4c). Thus, we found the CRRs of ribs could be a strong candidate of indicator for a good state of preservation of DNA in the skeletal remains.

**Figure 4.**
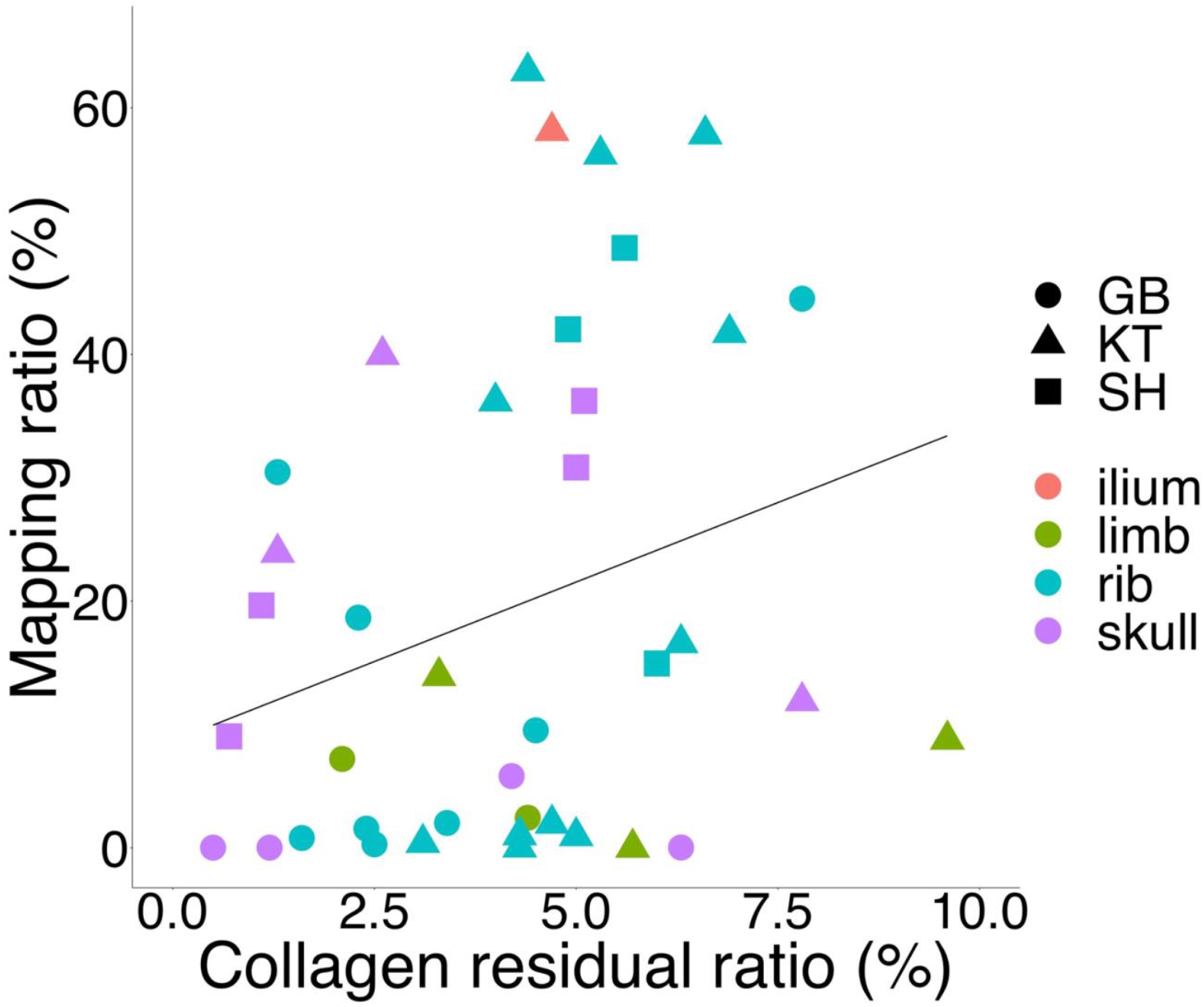
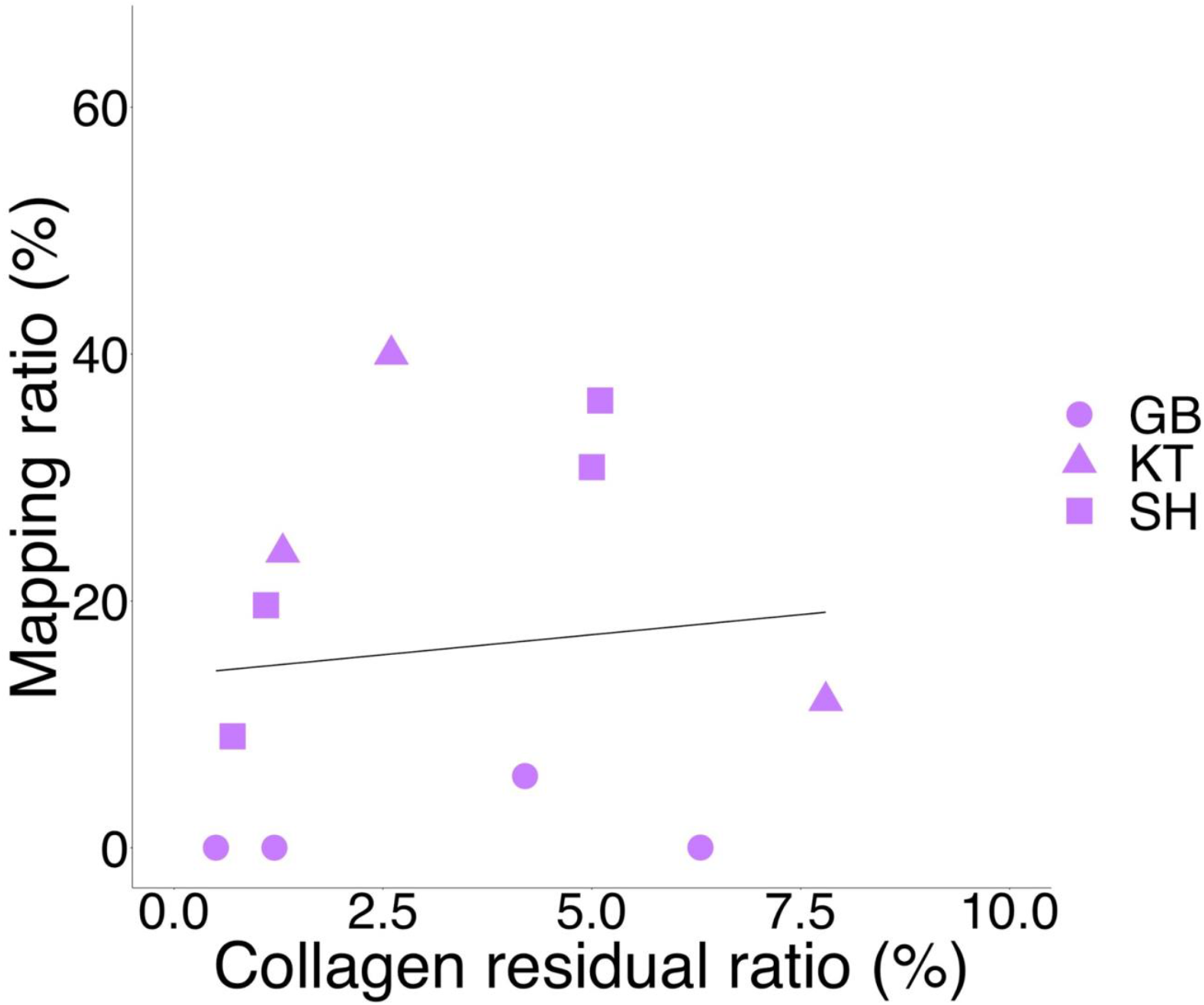
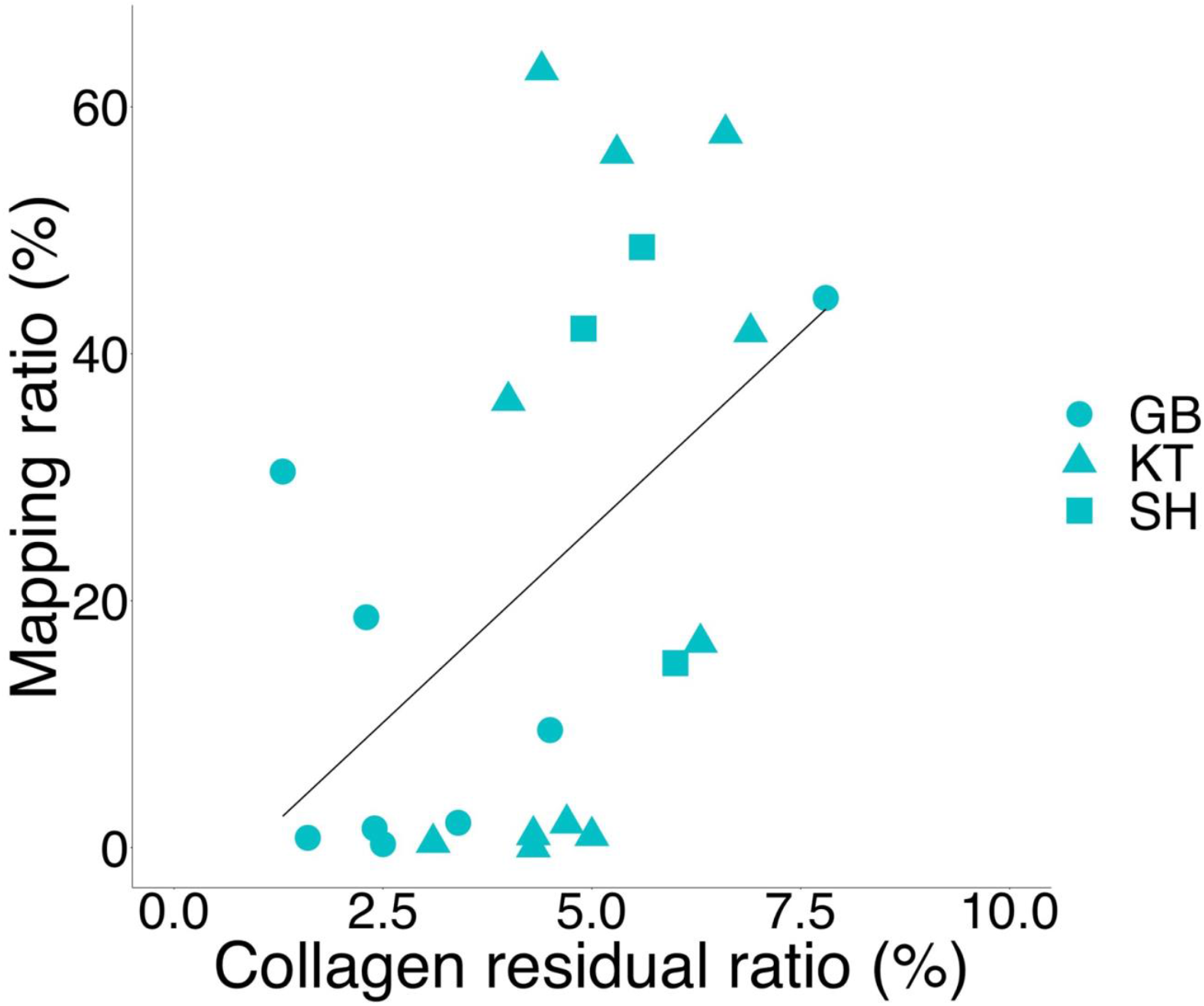
The association between the CRRs and the MRs. Circles, triangles, and squares represent the Gionbara shell-mound site, the Kikumatenaga shell-mound site, and the Saihiro shell-mound site, respectively, which include the bone parts, limb, ilium, rib, and skull, as shown. (a) The CRRs and the MRs in the three Jomon sites were shown. There was no association between them (*p* = 0.090). (b) The CRRs and the MRs in the skulls (n = 11) from the three Jomon sites were shown. There was no association between them (*p* = 0.745). (c) The CRRs and the MRs in the ribs (n = 22) from the three Jomon sites were shown. There was the association between them (*p* = 0.020).

## Discussion

In this study, we used the mapping ratios (MRs) based on the reads from NGS to assess the amount of endogenous DNA, instead of the conventional PCR amplification. As environmental factors, we tested in- or out-shell-stratum and the dating of the bones excavated from three archaeological sites (Figure 1 and Table 2). We found no association between the environmental factors and the MRs (Figures 2 and 3). As a molecular factor, we looked at the collagen residual ratio (CRR). When we tested combining all the samples, we had no association between the CRRs and the MRs. Similarly, we examined each archaeological site, we had no association between them (Figure 4a). Meanwhile, when we focused on the rib bones, we found a significant association between the CRRs and the MRs (Figure 4c). Though this might be an important finding, we still had a possibility of a spurious association because of combining the rib samples from three archaeological sites that have different preservation conditions. Therefore, we conducted a regression analysis adjusted for the site (i.e., sites were included in a linear model as dummy variables). The results revealed that the sites were not significantly associated with MR, although the association between MR and CR in ribs was marginally significant (Supplementary Table 1). Hence, we conclude that CRR in rib bones was significantly associated with MR.

It is still unclear the association between the environmental factors and DNA preservation. The result shown in Figure 1 that there was no association in DNA survival between in- and out-shell-stratums was unexpected, as soil pH should have a significant bearing on DNA preservation. Because the number of samples that could be compared in this study might be too small, additional analysis with a larger number of samples will be required in the future. On the other hand, as regards the lack of an association with calibrated radiocarbon dates shown in Figure 2, the result was to be expected. It is known that DNA is degraded soon after the death of the organisms and the length of DNA (less than 100 bp in mtDNA) does not change for a long period (Sawyer et al., 2012). The degradation of DNA could be approximated as exponential decay (Allentoft et al., 2012). Our results are thought reasonable, as the samples are more than 3,000 years postmortem, so a difference in age of several hundred years is not expected to affect DNA preservation.

Our study shows that a biomolecular factor, CRR, looks very promising as an indicator of DNA preservation. We found that the CRR of the ribs particularly seems to be suitable as the indicator. Because ribs are flat bones whose thick spongy bone is sandwiched between two layers of thin compact bones, the degradation of collagens in rib bones could be faster than the other hard bones, such as skulls, which might be more parallel to the rate of DNA degradation. Collagen is always extracted for dating which must be done in ancient genome analyses, so no additional bone-crushing for calculating CRRs is involved. It is very attractive that the CRR can be calculated by FTIR without crushing bones (Naito et al., 2020). Thus, further research may lead to the development of technology that enables the measurement of DNA preservation by non-crushing-bones.

## Acknowledgments

This study was supported by JSPS KAKENHI Grant-in-Aid for Scientific Research (A) to 18H03590 and 22H00020 to R.T.

## Figure legends

Supplementary Figure 1. Damage pattern of DNA molecules in libraries in this study. The x-axis means nucleotide position from the end of 5’ and 3’, and the y-axis means the frequency of substitution for reference human genome (hg19). The Blue line represents the C-to-T substitution and the red line represents the G-to-A substitution.

