## Supplementary Table 1 & Supplementary Figure 1 for "Collagen of ancient bones gives an indication of endogenous DNA preservation based on the next generation sequencing technology"

| | Regression coefficient (Standard error) | P-value |
|---|---|---|
| Collagen residual ratio of ribs | 5.732 (3.108) | 0.082 |
| KT[a] | 1.490 (11.190) | 0.896 |
| SH[a] | 8.693 (15.870) | 0.591 |

[a]GB was the reference.

Supplementary Figure. 1

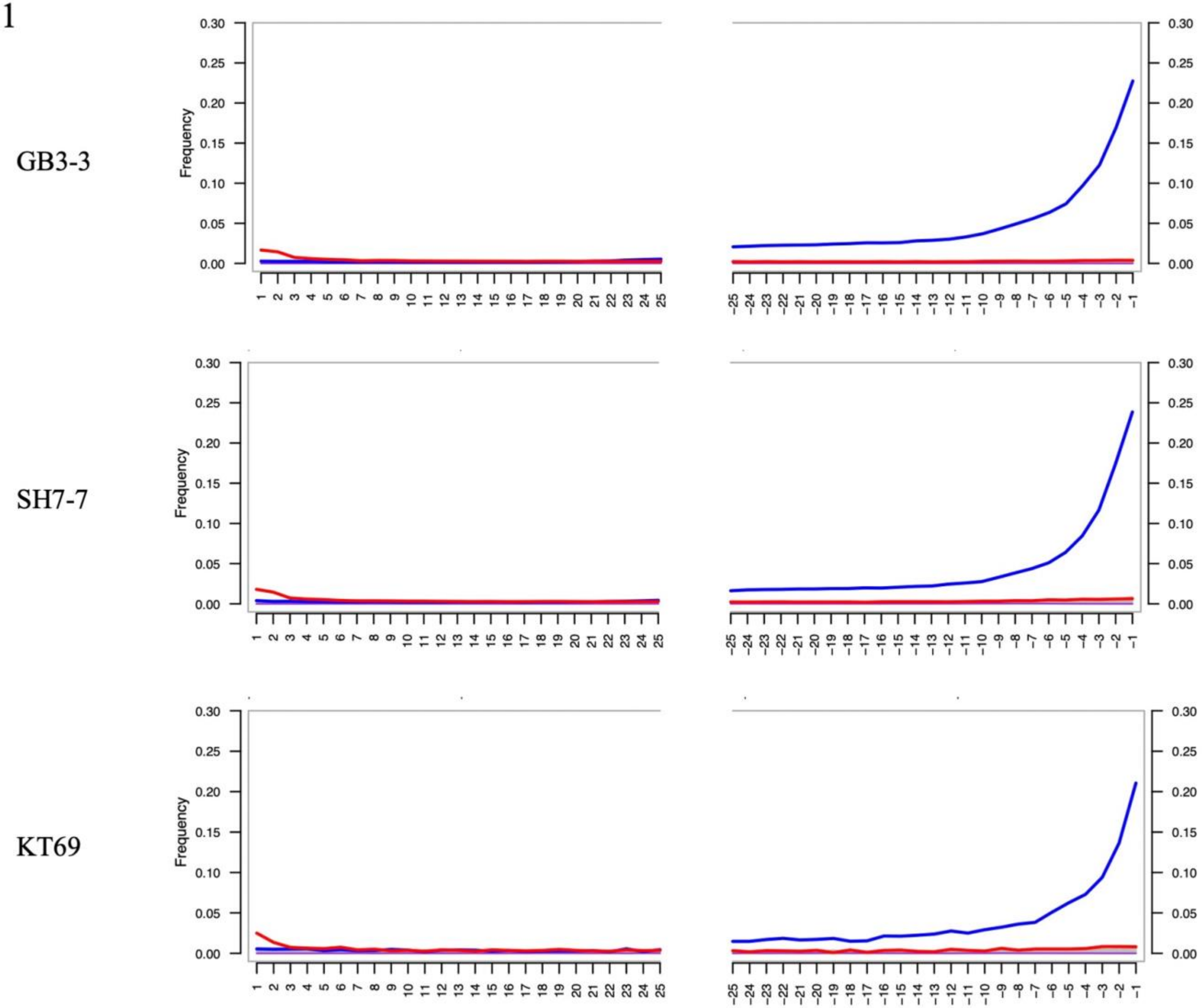
